# Rapid Speciation Characterized by Incomplete Lineage Sorting in the Globally Distributed Bacterium *Sulfitobacter*

**DOI:** 10.1101/2022.06.15.496264

**Authors:** Xiaojun Wang, Keigo Uematsu, Haruna Nakamura, Tetsuya Akita, Kei Kimura, Guangyu Li, Xingqin Lin, Dechao Zhang, Xiao-Hua Zhang, Chaomin Sun, Ingrid Obernosterer, Yuji Tomaru, Zongze Shao, Christian R Voolstra, Hideki Innan, Haiwei Luo

**Author notes:** These authors contribute equally to this work (Xiaojun Wang, Keigo Uematsu). **Corresponding author:** Hideki Innan Graduate University for Advanced Studies, SOKENDAI, Japan, Haiwei Luo School of Life Sciences, The Chinese University of Hong Kong Shatin, Hong Kong SAR China.

## Abstract

Incomplete lineage sorting (ILS), where ancestral polymorphisms persist through rapid speciation, is a hallmark of eukaryotic radiations but undocumented in prokaryotes. We report ILS associated speciation in the globally distributed marine bacterium *Sulfitobacter*, analyzing 74 strains from 16 oceanic regions and 15 ecosystem types. Genomic analysis resolved over ten genetically monomorphic clades sharing near uniform whole genome divergence (average nucleotide identity ≈95%). However, individual genes and even adjacent nucleotide sites support conflicting phylogenetic relationships among clades. The median length of nearly identical sequence blocks supporting alternative topologies is less than 50 base pairs, far shorter than typical adaptive cassettes. This within gene, site level discordance provides a diagnostic signature of ancestral polymorphism retention following rapid fragmentation, ruling out adaptive gene sweeps. Demographic inference places this radiation on a timescale of tens of millennia, 100 to 1,000 fold faster than eukaryotic ILS, and traces the clades to a hyper polymorphic, highly recombining ancestor. Neutral simulations show that transient mutation surges or population bottlenecks alone can generate this pattern, without adaptive selection or ecological partitioning. By bridging eukaryotic and prokaryotic speciation paradigms, our findings demonstrate that ILS, previously considered exclusive to sexual eukaryotes, can drive rapid, neutral diversification in bacteria, expanding the mechanisms known to generate prokaryotic biodiversity.

## Introduction

The evolutionary histories of genes within species often conflict due to processes such as incomplete lineage sorting (ILS), where ancestral polymorphisms persist through successive speciation events. ILS was first formalized theoretically by Tajima (1983) ^1^ and Pamilo & Nei (1988) ^2^, and subsequently gained wide empirical recognition through studies of human, chimpanzee, and gorilla genomes, where different genes supported conflicting evolutionary relationships ^3^. The phenomenon arises when rapid population divergences outpace the sorting of ancestral genetic variation, creating incongruent gene trees that obscure phylogenetic relationships ^2^. ILS is not merely a source of phylogenetic discordance; it can also lead to genomic regions following alternative evolutionary trajectories. For example, in hominid primates, approximately 15% of the human genome is more closely related to gorilla than chimpanzee, despite chimpanzees being humans’ closest relatives, a signature of ancestral polymorphism retention during rapid divergence ^4,5^. ILS has been documented across all major eukaryotic lineages ^6–12^, including plants ^13–16^, fungi ^17,18^, and unicellular protists ^19,20^, and it is particularly prevalent in groups undergoing evolutionary radiations ^21^. Its role in prokaryotes has remained entirely unexplored.

In eukaryotes, a large ancestral population and short time intervals between successive speciation events together drive ILS ^2^. Because population sizes of prokaryotic species are likely many orders of magnitude greater than that of multicellular eukaryotes, ILS would be expected to have an even stronger influence on prokaryotic phylogenomic patterns, assuming a roughly constant rate of species diversification through time ^22^. Nevertheless, the phenomenon has never been documented in prokaryotes, raising the question of why it has been overlooked.

Prokaryotic speciation research has long been guided by two models that emphasize natural selection. The Biological Species Concept (BSC) ^23,24^, adapted from eukaryotes ^25,26^, posits that speciation involves reduced gene flow between incipient species, although it permits ongoing cross-species genetic exchange unlike its eukaryotic counterpart ^24^. The Ecotype Model ^27^, in contrast, emphasizes ecological divergence as the primary driver, where niche adaptation promotes genetic isolation. Both frameworks assume that natural selection is essential for speciation, acting either to reduce recombination (BSC) or to exploit new niches (Ecotype), whereas neutral processes such as genetic drift are dismissed as insufficient in bacteria because of their large population sizes ^28,29^. Because ILS is a neutral process (or at least does not require adaptive selection), it was never considered as a possible mechanism of bacterial diversification. This historical bias explains the gap between theoretical applicability and empirical documentation.

Here, we studied *Sulfitobacter*, a globally distributed marine bacterium, using phylogenomic analysis of 74 strains sampled from diverse oceans and ecosystems. We uncovered a pattern of rapid speciation characterized by persistent phylogenetic discordance even at the level of individual genes and adjacent nucleotide sites. This signature, together with demographic inference and simulations, indicates that these clades arose from a highly polymorphic, recombining ancestral population through ILS, without requiring adaptive selection or ecological niche partitioning. Our findings thus bridge eukaryotic and prokaryotic speciation paradigms, demonstrating that even neutral processes can drive rapid bacterial diversification.

## Results

### A globally distributed Sulfitobacter lineage comprises multiple equidistant, genetically monomorphic clades

We analyzed 74 *Sulfitobacter* strains collected from 16 oceanic regions and 15 ecosystem types (Fig. 1), including 70 newly isolated genomes from diverse marine environments and four public genomes (Table S1, Fig. S1). Phylogenomic reconstruction resolved these strains into 14 clades. Nine clades were singletons (each represented by a single strain), one clade (C6) contained two strains, and four clades (C1, C2, C9, C12) each contained three or more strains (Fig. 1). To exclude potential biases from clonal expansion during sample handling, we retained 45 non-duplicate strains (identical clade, ecosystem, location, and cruise) for all subsequent analyses.

**Fig. 1.**
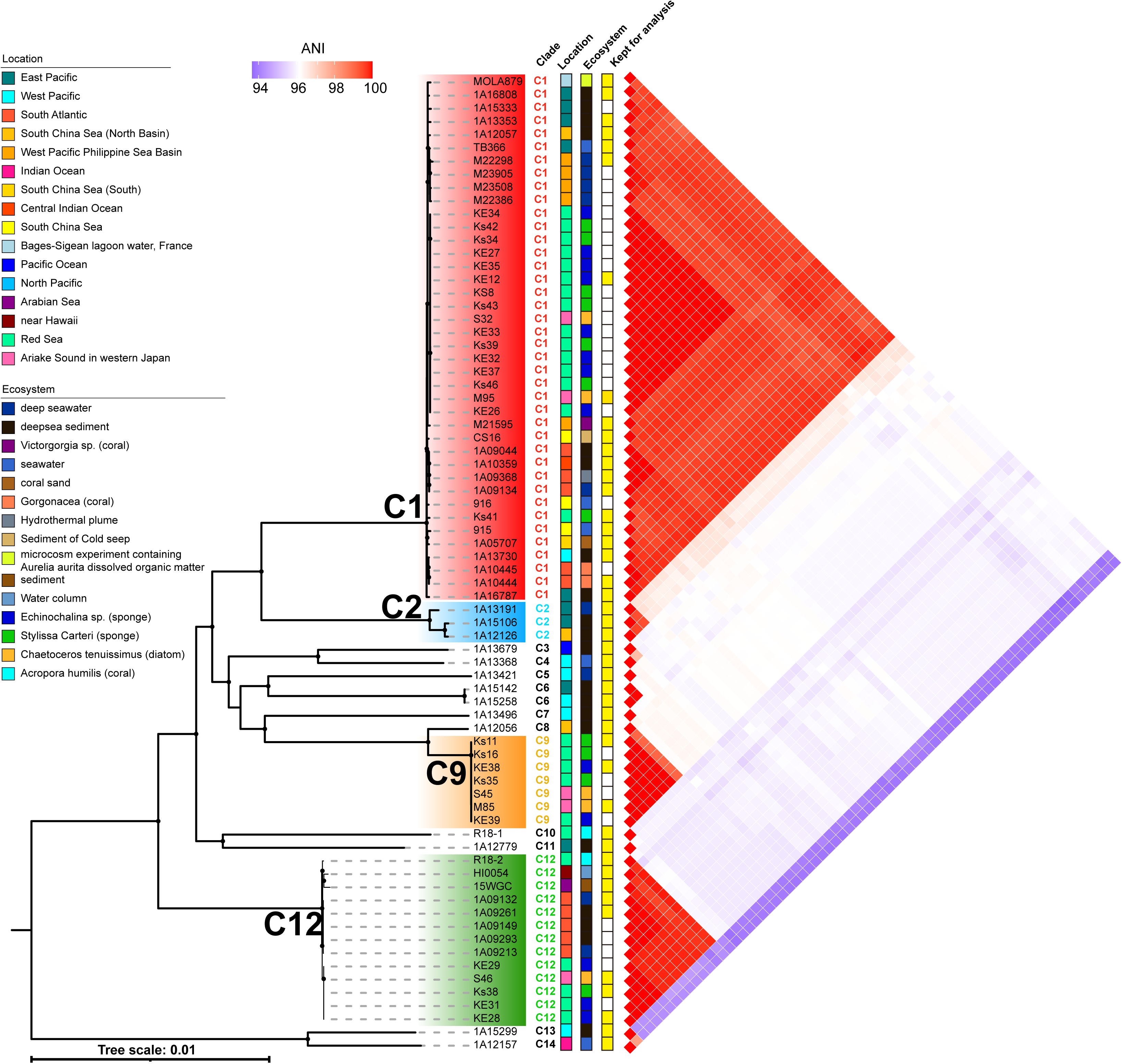
Phylogenomic tree and wholeDgenome average nucleotide identity (ANI) of 74 *Sulfitobacter* strains. (Left) The IQ-TREE maximum-likelihood phylogenomic tree of 74 *Sulfitobacter* strains, which is a subset of a comprehensive phylogenomic tree of *Sulfitobacter* (Fig. S1). Internal clades are labeled by ID, and clades with at least three available genomes are highlighted in color. Sampling source, including geographic location and ecosystem type, is indicated for each strain. Each clade is marked with an ID on its internal branch, and those clades each with at least three genomes available are marked with color. For each strain, the sampling source (geographic location and ecosystem type) is provided. If multiple strains were isolated from the same sample source, only one representative strain was retained for further analysis and is denoted as a yellow rectangle. (Right) The heatmap of the whole-genome ANI of the 74 strains.

The four multi-member clades (C1, C2, C9, C12) exhibited extreme genetic monomorphism. Within-clade average nucleotide identity (ANI) ranged from 99.64% to 99.99% (Fig. 1), and single nucleotide polymorphism (SNP) densities spanned 3 to 4,027 SNPs per Mbp (Table S2). This near identity persisted even when strains originated from markedly different habitats. Clade C1, for example, included isolates from coral, deep sea sediment, and hydrothermal plume sources across the South China Sea, Indian Ocean, and West Pacific, yet maintained 99.68% ANI. Clade C12, with strains from sponge, diatom, and coral hosts across distant geographic locations, exceeded 99.86% ANI (Fig. 1).

Between clades, pairwise ANI values were uniformly close to 95%, the conventional prokaryotic species threshold. This uniformity was independent of phylogenetic relatedness inferred from the concatenated tree (Fig. 1). In that tree, C1 was more closely related to C9 than to C12, but the ANI values were nearly identical (96.13% versus 95.96%). Only two clade pairs (C8/C9 and C13/C14) showed higher ANI (above 97%).

### Genome-wide phylogenetic incongruence between and within orthologous gene families

We assessed phylogenetic incongruence across the genome by analyzing 2,089 single copy orthologous gene families shared by the 45 non duplicate strains, each containing more than three non-redundant gene sequences for tree construction. Because evaluating all possible phylogenetic trees for 14 clades is computationally intractable (nearly 3.162×10^11^ unrooted trees), we used quartet analysis. A quartet consisted of four strains from four distinct clades, with three possible unrooted topologies describing their relationships (Fig. 2A). We aggregated results across quartets to infer genome-wide patterns of discordance.

**Fig. 2.**
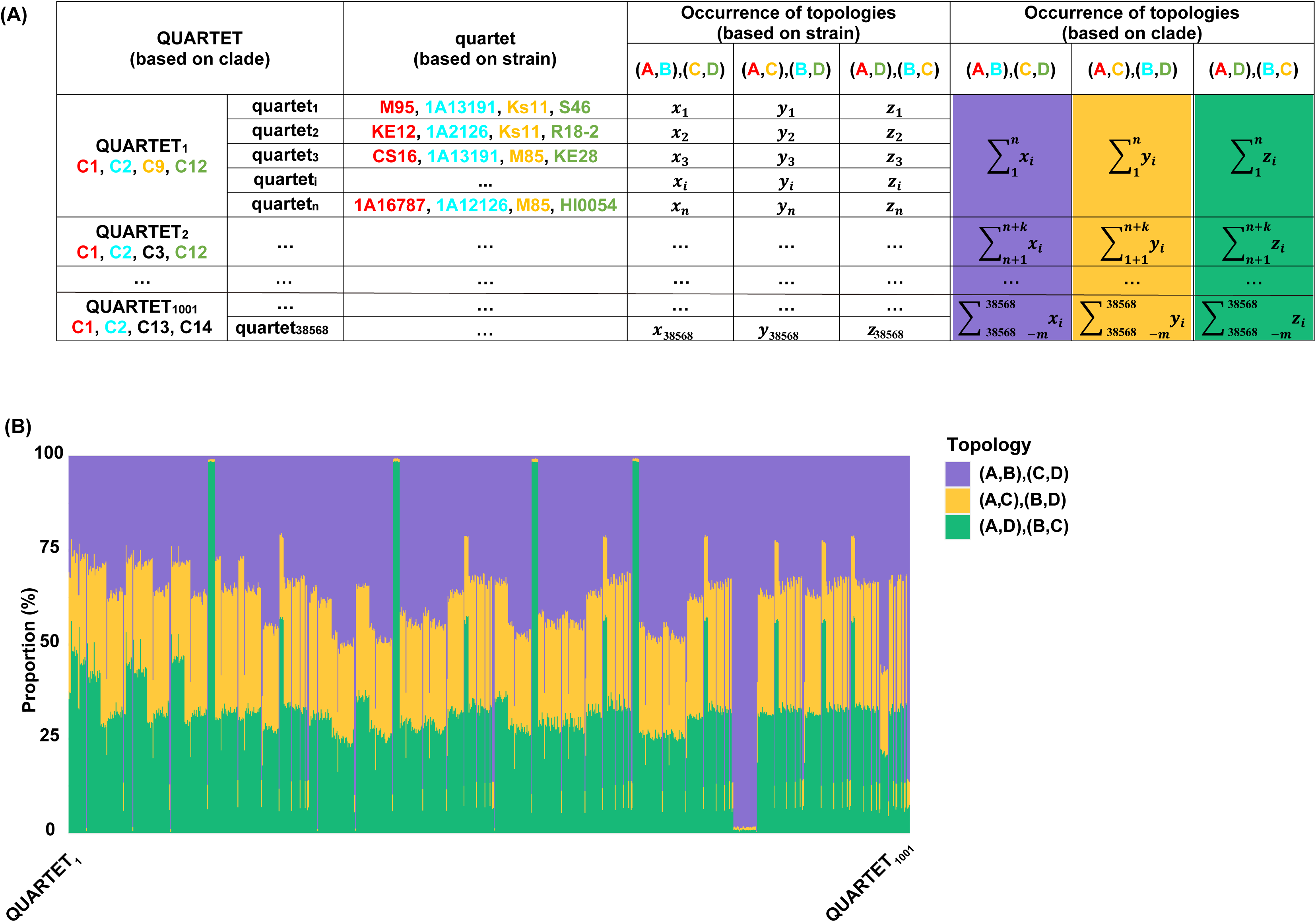
Quartet analysis for detecting phylogenetic incongruence. (A) There are a total of 38,568 quartets, each comprising four strains sampled from four different clades. These quartets are categorized into 1,001 distinct groups (QUARTETs), each consisting of members from the same combination of four different clades. For each quartet, the occurrence of each possible topology was determined with QuartetScores across 2,089 unrooted gene trees. These occurrences were then summed across all quartets within a QUARTET to calculate the total occurrence of each topology for that QUARTET. (B) The proportion of the 2,089 unrooted gene trees supporting different quartet topologies for each of the 1,001 QUARTETs, based on the columns with colored backgrounds in panel (A).

We examined 38,568 quartets composed of strains from four different clades, and grouped them into 1,001 “QUARTETs” based on their clade composition (e.g., QUARTET_C1-C2-C9-C12; Fig. 2A). For each QUARTET, we calculated the proportion of gene trees supporting each of the three topologies. In 87% of QUARTETs, no single topology dominated; gene trees were split nearly equally among all three possibilities (Fig. 2B). For QUARTET_C1-C2-C9-C12, no topology received support from more than 44% of genes (Fig. 2B). Only quartets involving very closely related clades (e.g., C8/C9, ANI ≈ 99.05%) showed a strong bias toward one topology, consistent with their recent divergence.

Incongruence also appeared at the level of individual nucleotide sites. We identified 113,206 phylogenetically informative SNPs (phylo-SNPs) within the 2,117 core genes shared by 45 non duplicate strains. These phylo-SNPs were defined as sites conserved within each clade but variable between clades. In a representative gene (Fig. 3A), seven distinct types of biallelic phylo-SNPs partitioned the four focal clades (C1, C2, C9, C12) into different groupings. Four phylo-SNP types separated one clade from the other three (e.g., C1-specific alleles), whereas three types divided the clades into pairs (e.g., C1+C2 vs. C9+C12). The incompatibility between pairwise SNPs has been used as a signature of recombination breakpoints ^30^.

**Fig. 3.**
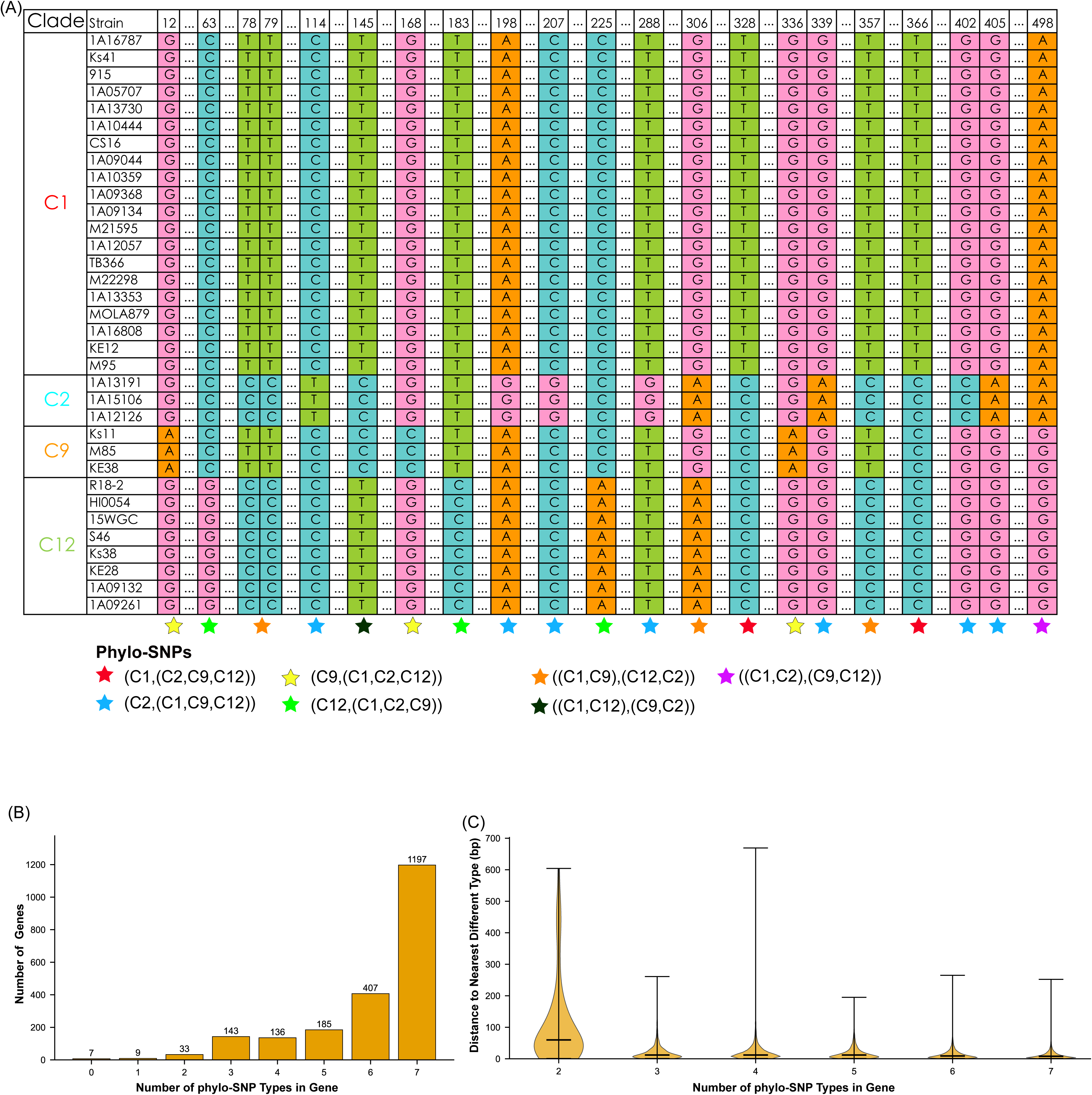
PhyloDSNP patterns within a representative core gene family and distribution of phyloDSNP types across all core genes. (A) An example of the phylo-SNP patterns derived from segments of a core gene family shared by the four focal clades (C1, C2, C9, and C12). Strains are ordered by clade affiliation. Four clades can have a maximum of seven possible biallelic phylo-SNPs, which are all found in this example and marked with colored stars based on the tree topology they support. Conserved nucleotide sequences between phylo-SNPs are represented with ellipsis. (B) A total of 2,117 single-copy orthologous gene families were classified based on the number of phylo-SNP types. The bar plot shows the number of gene families containing each possible number of phylo SNP types (from 0 to 7). SNPs that are not phylo-SNPs are not considered here. (C) The distance (bp) between the nearest phylo-SNPs of conflicting topology within individual genes.

Nearly 99% of the 2,117 gene families contained at least two types of phylo-SNPs, and more than half harbored all seven phylo-SNP types (Fig. 3B). To measure the physical length of nearly identical regions between clades, we calculated the distance (bp) between the two closest phylo-SNPs of different types within each gene. Across all gene families, these distances were short, with a median of less than 50 bp regardless of the number of phylo-SNP types present (Fig. 3C).

### Recent recombination is rare between clades

We conducted a second quartet analysis to quantify recent recombination events between clades. Unlike the previous analysis that assessed genome-wide incongruence among four different clades, this analysis focused on quartets composed of strains from either two or three clades (Fig. 4A). In such quartets, a gene tree where strains from the same clade are separated into different branches would indicate recent homologous recombination between clades (Fig. 4B). A total of 105,512 quartets were analyzed and grouped into 400 QUARTETs based on their clade composition (Fig. 4A). Of all gene trees, 1.25% supported fully mixed clade membership (Fig. 4C). Among these, 72% involved clades C8 and C9, which shared an ANI of 99.05%. After excluding quartets involving C8 and C9, the proportion of gene trees with mixed clade membership was 0.35%. We also examined homoplastic SNPs. Of 119,143 biallelic polymorphic sites (BiPs), only 3,086 (2.6%) were homoplastic, and 77% of these homoplastic SNPs clustered in 46 genes. We scanned for regions of high nucleotide identity (greater than 99%) between clades using sliding windows of 1,000 bp with a 500 bp step. For the C1/C2 pair, 489 of 5,064 windows (9.6%) showed high identity. For all other clade pairs, the number of high-identity windows ranged from 19 to 32 out of 5,064 (less than 0.6%) (Fig. S2). The extremely low nucleotide diversity within each clade (Table S2) prevented reliable detection of recombination events within clades, which precluded a direct rate comparison between within clade and between clade recombination.

**Fig. 4.**
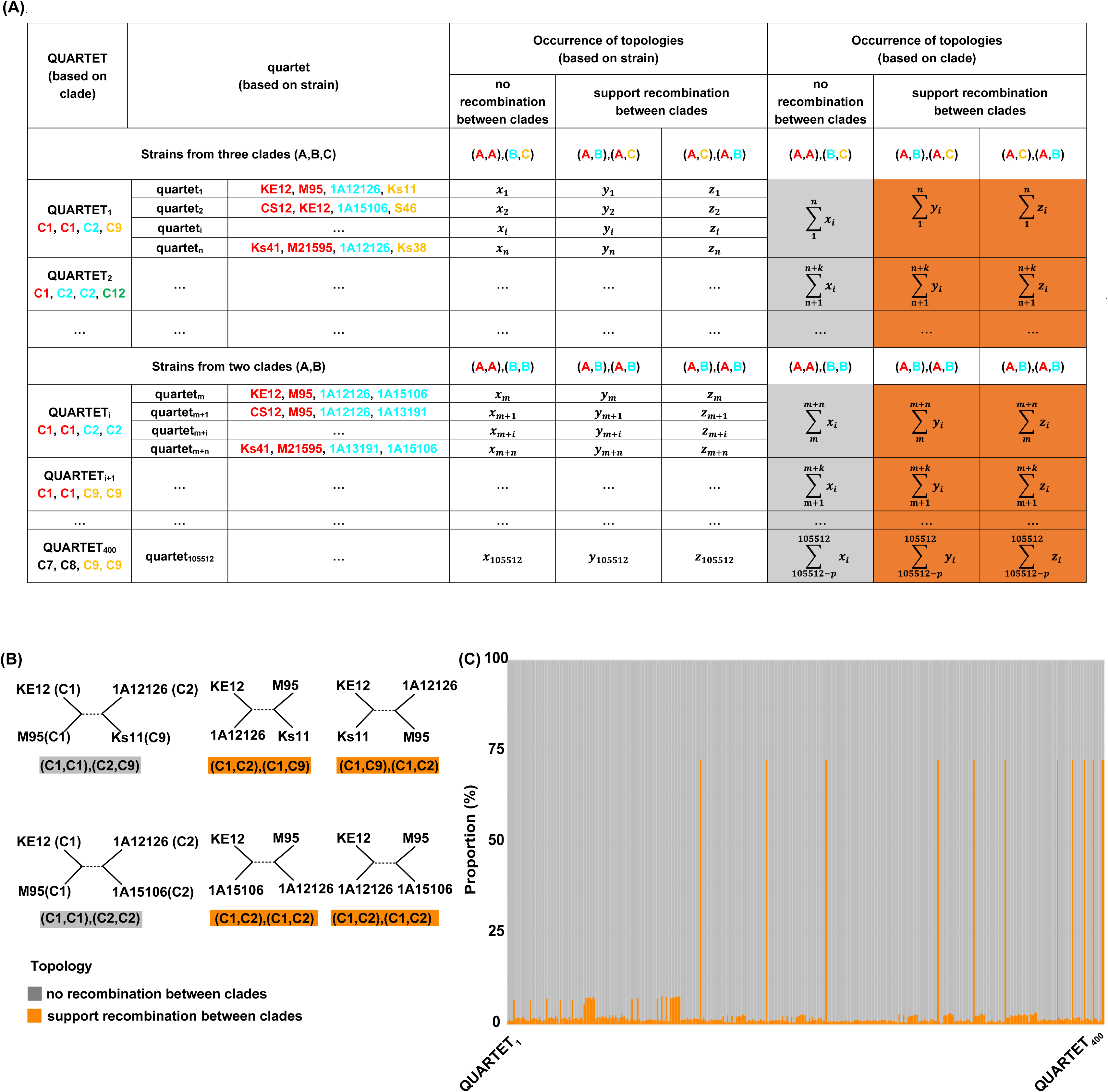
Quartet analysis for detecting recent recombination between clades. (A) There are a total of 105,512 quartets, each comprising four strains that involve either two or three different clades. These quartets are grouped into 400 quartet groups (QUARTETs), each consisting of members from the same combination of clades. The method for calculating the support for each topology is identical to that shown in Fig. 2A. (B) Three possible quartet topologies of two example quartets. Four strains are from either three or two different clades. The quartet topology where same-clade members cluster together (highlighted in gray) indicates no signal of recent recombination between clades. Conversely, the quartet topologies where same-clade members are separated into different clades in the gene tree (highlighted in orange) support homologous recombination between clades. (C) The proportion of the 2,089 unrooted gene trees showing evidence for, or against, recent recombination between clades for each of the 400 QUARTETs.

### Limited ecological, genetic, or biogeographic barriers between clades

We identified clade specific and clade exclusive gene families from the accessory genome (Table S4). For clade C9, a set of four orthologous gene families involved in polyhydroxyalkanoate (PHA) synthesis was present, whereas other clades also carried genes with the same functional annotation but at lower copy numbers (Table S5). No other clade possessed a complete metabolic pathway absent from the others (SI Text S1.1). We examined the distribution of microbial defense systems, including restriction modification (R M) systems and CRISPR Cas systems (Fig. S3). Copy numbers of type I to IV R M systems varied among genomes, but no clade specific pattern emerged. No CRISPR Cas system was detected in any of the sequenced genomes.

We also assessed geographic distributions of the four focal clades. Multi member clades each included strains isolated from distinct ocean basins (Fig. 1), indicating that no clade was restricted to a single geographic region.

### Limited adaptive sweep signatures in core and accessory genomes

We examined the core genome for signatures of adaptive sweeps by analyzing 2,117 single copy orthologous gene families. Among these, 360 gene families exhibited at least 99% mean pairwise nucleotide identity between at least one clade pair. Of these 360 families, 205 (57%) showed high identity only between the sister clades C1 and C2 (Table S6). The remaining 155 families showed high identity between at least one non sister clade pair. In these 155 families, the dominant functional category was ribosomal proteins and translation machinery (COG J, accounting for 25% of the families). Within the 155 families, nine gene clusters were identified, defined as at least two contiguous loci in the genome. Five of these clusters corresponded exclusively to ribosomal proteins, whereas the remaining four either lacked functional annotation or could not be assigned to any well defined pathway (Table S6).

We also tested whether accessory genes shared among subsets of clades but absent from others have been homogenized by adaptive sweeps. For each such subset specific gene family, we compared its between clade nucleotide identity to the genomic background identity for the same clade pair. Across six testable clade pairs, subset specific genes showed equal or lower identity than the background (Mann Whitney U test, all *P* > 0.05; Fig. S4). Only five gene families exceeded 99% between clade identity, of which four involved the sister clades C1 and C2 (Table S4).

### Demographic inference reveals rapid successive speciation events

We analyzed linkage disequilibrium (LD) decay patterns across the genome to characterize historical recombination rates. Pairwise LD (*r²*) was calculated for 113,206 biallelic phylo-SNPs. LD decayed to background levels within 500 bp (red dots in Fig. 5A), matching the expectation for unlinked biallelic phylo-SNPs (horizontal dashed line in Fig. 5A). This rapid LD decay indicated that recombination had shuffled alleles sufficiently that distinct genomic regions could have independent coalescent histories. We therefore inferred the demographic history of the four focal clades (C1, C2, C9, C12) using site frequency spectrum (SFS) analysis. The full dataset of 14 clades would have provided broader phylogenetic resolution, but the SFS becomes increasingly sparse with more populations, reducing statistical power. We focused on the four clades to ensure robust parameter estimation. Using dadi.CUDA ^31^, we modeled a four-population divergence scenario (Fig. 5B). Assuming a mutation rate of 10^-10^ substitutions per site per generation, consistent with *Sulfitobacter pontiacus* EE-36 ^ref32^, we obtained an ancestral population size (*N_0_*) of 2.08 × 10^8^. The model yielded three sequential splits within approximately 2.26% of *N_0_* generations. Each split corresponded to a sharp reduction in effective population size (*N_e_*). The inferred speciation order was ((C1, C2), C9), C12), which matched the topology of the concatenated phylogenomic tree (Fig. 1). To estimate the absolute timescale of the radiation, we assumed a generation time of one day, as previously determined for pelagic *Roseobacter* ^33,34^. Under this assumption, the entire radiation occurred within approximately 20,000 years.

**Fig. 5.**
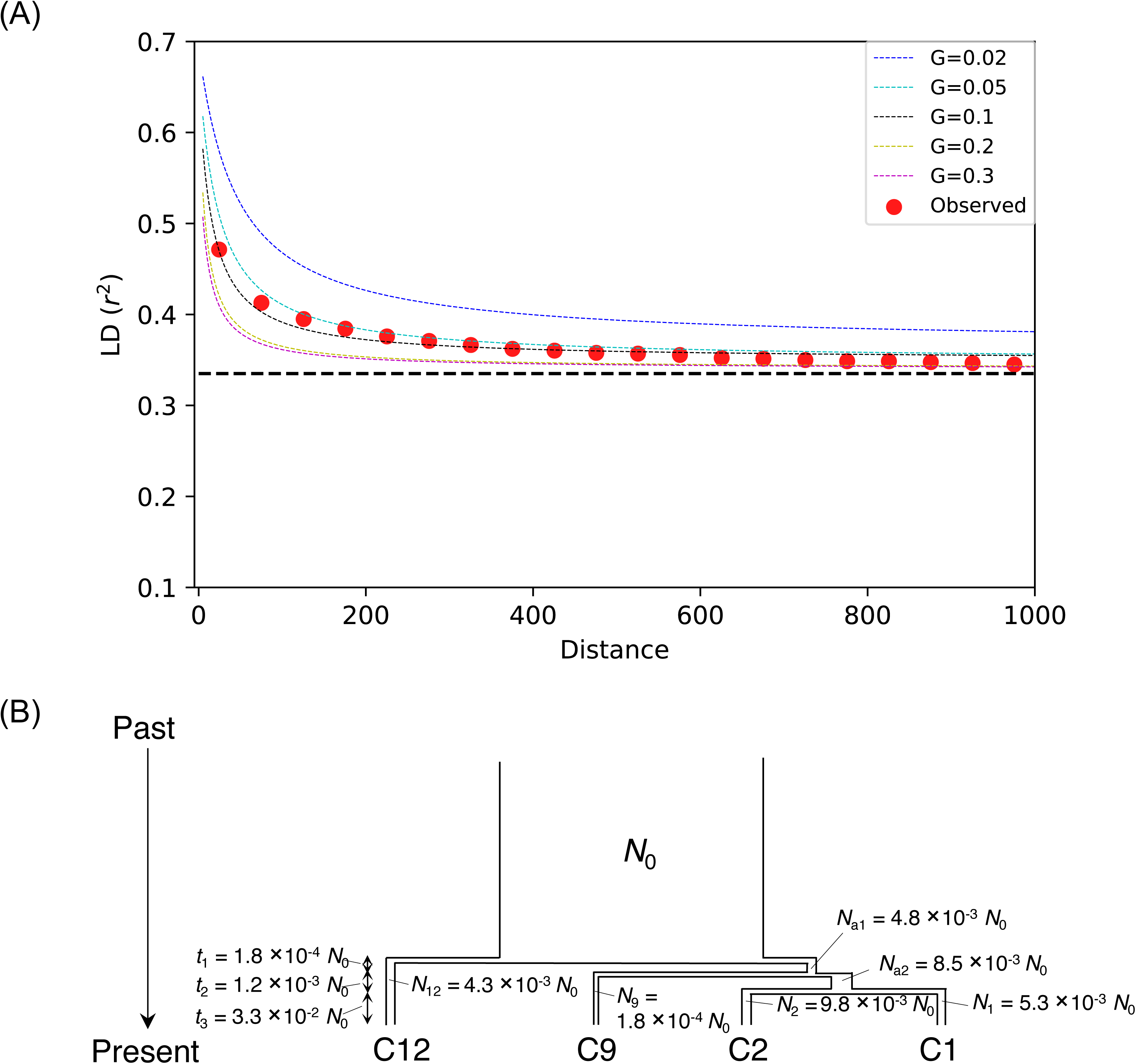
Linkage disequilibrium (LD) decay and demographic model for the four focal clades (C1, C2, C9, and C12). (A) The decay of LD across the four focal clades. The horizontal and vertical axes represent the distance (bp) between pairwise biallelic phylo-SNPs and LD based on the *r^2^*measure, respectively. The horizontal dashed line marks the expected LD under the inferred demography when the distance of pairwise biallelic phylo-SNPs is ∞. Distances are binned with a window size of 50 bp, and the red dots denote the LD values calculated from the *Sulfitobacter* population under study. The dashed curves represent the decay of LD under different population recombination rates (*G*), assuming the demographic model shown in (B) and a recombination tract length of 1,000 bp. (B) Inferred demographic history of the four focal clades. The model assumes an ancestral population of size *N_0_*that undergoes three sequential splits. The size of each descendant population and the duration between speciation events are shown relative to *N_0_*. Model parameters were inferred using dadi.CUDA.

### High recombination rate in the ancestral population

We quantified the ancestral recombination rate (*G_0_*) of the four focal clades by performing simulations under varying *G_0_* to fit the observed pairwise LD (*r*^2^) for 113,206 biallelic phylo-SNPs. The simulations identified *G_0_* = 0.1 as the best fit to the observed LD decay (Fig. 5A). This result was robust to adjustments in the average recombination tract length *L* (Fig. S5). We then calculated the ancestral population mutation rate (θ*_0_***=** 0.041) using the mutation rate of *Sulfitobacter pontiacus* EE-36 (10^-10^ substitutions per site per generation) and the inferred ancestral population size (*N_0_*= 2.08 × 10^8^). The resulting recombination-to-mutation rate ratio was 2.44.

### Neutral demographic shifts generate rapid diversification in simulations

We performed individual-based simulations that explicitly excluded adaptive selection to test whether neutral demographic processes alone could produce patterns similar to the observed *Sulfitobacter* radiation. The null model assumed a haploid ancestral population divided into 10 subpopulations, each with 1,000 individuals, 10 loci, a recombination rate of 0.05 per generation, a mutation rate (μ) of 10^-6^ substitutions per locus per generation, and a migration rate of 10^-5^ events per generation. Under these baseline conditions, only one new lineage appeared within 2 × 10^7^ generations (SI Text 1.2, Fig. S6). We then simulated two neutral perturbations. In the first, we increased μ 100-fold (to 10^-4^) for a short period (within 5 × 10^6^ generations) in the middle of the simulation. This transient mutation surge generated a rapid accumulation of haplotypes, followed by population fragmentation into multiple isolated lineages (Fig. 6A). After returning μ to the baseline value, descendant clades remained genetically isolated despite the continued high migration rate. In the second perturbation, we reduced each subpopulation size to 100 individuals (10% of its ancestral value) for the same duration (5 × 10^6^ generations). This bottleneck also fragmented the population into many isolated lineages (Fig. 6B), with genetic divergence between subpopulations increasing sharply. After restoring the population size to 1,000 individuals, gene flow between clades did not resume.

**Fig. 6.**
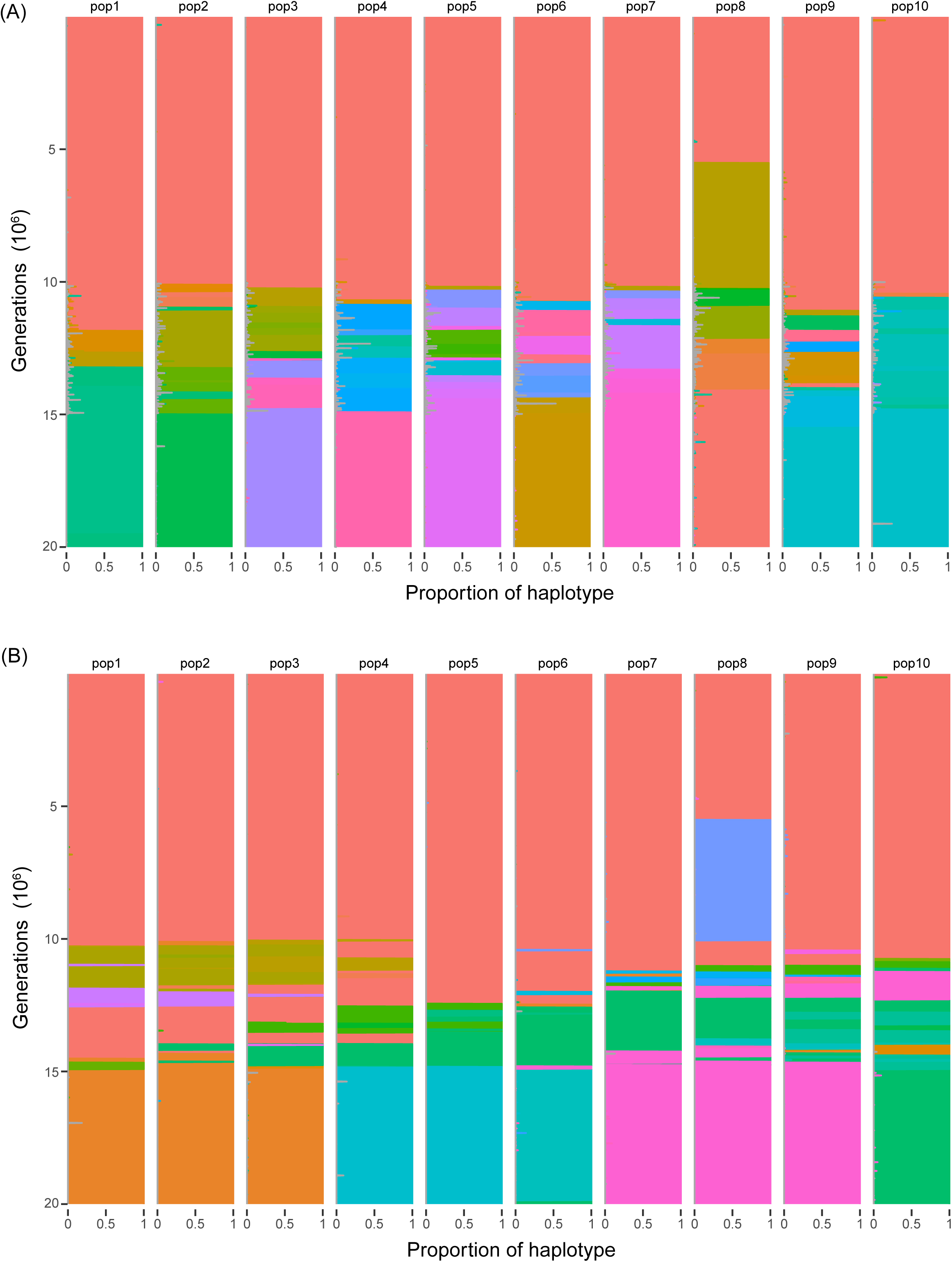
Neutral simulations of rapid diversification under a mutation surge and a population bottleneck. (A) Representative result from a neutral simulation under a surge of mutation rate. All parameters are the same as the null model (number of subpopulation *K* = 10, the number of haploid individuals in each subpopulation *N* = 1000, the number of loci *L* = 10, the recombination rate *p* = 0.05, the mean number of loci transferred in a single recombination event *r* = 1, mutation rate μ = 10^-6^, the strength of purifying selection *s* = 0.2, and migration rate *m* = 10^-5^), except that μ was increased to 10^-4^ between *t*=1.0×10^7^ and 1.5×10^7^ generations. (B) Representative result from a neutral simulation under a bottleneck mode. All parameters are the same as the null model, except that *N* was reduced to one-tenth between *t*=1.0×10^7^ and 1.5×10^7^ generations.

## Discussion

### Parallels and differences with eukaryotic incomplete lineage sorting

The *Sulfitobacter* clades exhibit pervasive phylogenetic discordance even at the level of individual genes and adjacent nucleotide sites, with nearly identical sequence blocks between phylo-SNPs supporting alternative topologies having a median length of less than 50 base pairs. This pattern, together with uniform between clade divergence (≈95% ANI) uncorrelated with evolutionary distance, indicates incomplete lineage sorting (ILS) following rapid fragmentation of a highly polymorphic ancestral population.

The *Sulfitobacter* radiation mirrors ILS patterns in eukaryotes, such as hominids and cichlids, where rapid divergence preserves ancestral polymorphisms. In both domains, incomplete sorting of ancestral variation generates genome wide genealogical discordance. However, the prokaryotic process diverges in tempo and underlying mechanism. Eukaryotic ILS typically unfolds over millions of years ^3,6–8^, whereas the *Sulfitobacter* clades achieved similar genomic discordance on a timescale of tens of millennia. This acceleration likely stems from three factors. First, a large ancestral population size (*N_e_* ≈10^8^) preserved extensive polymorphism by buffering against stochastic loss of genetic diversity through drift. Second, a short generation time (approximately one day) compressed mutation and recombination events into a fraction of eukaryotic timescales; this estimate is based on pelagic Roseobacter ^33,34^ and serves as a proxy that may not capture species specific variation but is consistent with related marine bacteria. Third, frequent homologous recombination for a bacterium (G_0_/θ_0_ = 2.44) distributed ancestral alleles across the population, creating a genomic mosaic where different loci carry independent coalescent histories. Although bacterial recombination rates are in general much lower than those in sexual eukaryotes ^35^, this combination of large population size, short generation time, and elevated recombination rate allowed ancestral polymorphisms to persist and sort incompletely during rapid radiation. The demographic inference and absolute timescale also assume a mutation rate of 10^-10^ substitutions per site per generation derived from a single *Sulfitobacter* strain ^32^; any variation in this rate would rescale the estimated times but does not alter the conclusion of rapid, ILS driven diversification.

### Adaptive gene sweeps do not explain the observed phylogenetic discordance

An alternative interpretation for pervasive phylogenetic incongruence is that adaptive gene sweeps spread nearly identical sequences across divergent lineages through horizontal transfer ^36^. Under this model, a generally adaptive mutation or gene set sweeps within one ecotype and then transfers into other ecotypes, driving sequential selective sweeps. The transferred segment would support a grouping of donor and recipient lineages, whereas the rest of the genome remains divergent between the lineages, generating genome wide incongruence. In *Sulfitobacter*, however, the length of shared nearly identical regions argues against this mechanism. An adaptive segment transferred as a functional unit would produce long, contiguous blocks of near identity across divergent clades. Instead, the nearly identical sequence blocks between phylo-SNPs supporting alternative topologies have a median length of less than 50 bp, and 99% of orthologous genes contain at least two conflicting phylo SNP types (Fig. 3B, C). Even within individual genes, different nucleotide sites support conflicting phylogenetic groupings. This pattern is inconsistent with the transfer of intact gene sized adaptive cassettes. Recombination erosion after transfer could theoretically shorten blocks, but that would leave some genes intact, especially those transferred most recently. The observation that nearly every gene harbors multiple conflicting phylo-SNP types contradicts such an erosion scenario. The data thus support ILS rather than adaptive sweeps as the primary cause of genealogical discordance.

### Classical bacterial speciation models do not account for Sulfitobacter diversification

The *Sulfitobacter* clades defy the prokaryotic Biological Species Concept (BSC). Under the BSC, incipient bacterial species experience reduced but persistent gene flow ^37–39^. In *Sulfitobacter*, however, recent recombination between clades is minimal (0.35% of gene trees after excluding C8/C9), approaching complete cessation. This pattern aligns more closely with eukaryotic-style reproductive isolation than with the prokaryotic BSC expectation of ongoing genetic exchange.

The Ecotype Model posits that bacterial diversity arises from ecological divergence, with periodic selection purging genetic diversity within ecotypes ^36,40^. A central prediction of this model is that genomic signatures of adaptive sweeps should be detectable and linked to ecological adaptation. In the core genome, the majority of high identity regions were confined to sister clades, and the few high identity regions between non sister clades corresponded to housekeeping operons rather than ecologically adaptive modules. No multi gene adaptive islands were detected. In the accessory genome, genes shared among subsets of clades did not show elevated between clade identity relative to the genomic background, and no such gene family was associated with an obvious ecological function. The absence of ecologically coherent sweep signatures indicates that genomic differentiation of *Sulfitobacter* clades does not conform to the Ecotype Model. We do not rule out the possibility that ecological differences have accumulated after divergence, but the observed gene content differences can also be explained by post divergence drift and differential gene loss rather than ecological partitioning that initiated speciation.

### Neutral demographic shifts can drive rapid diversification

The individual based simulations demonstrated that rapid speciation can occur in the absence of adaptive selection. Under baseline conditions (large population, moderate migration, low mutation), only one new lineage appeared over the simulated timescale. In contrast, a transient 100 fold surge in mutation rate or a temporary 90% population bottleneck generated rapid lineage diversification, producing multiple isolated clades within a short period. After the perturbation ended, gene flow between clades did not resume despite the return of high migration rates. These results demonstrate that neutral demographic shifts alone are sufficient to produce a pattern of rapid fragmentation and persistent isolation similar to that observed in *Sulfitobacter*. The simulations thus provide a proof of principle that selection is not required for bacterial speciation of this type. Mutation surges can generate the standing ancestral polymorphism necessary for ILS, whereas bottlenecks can intensify drift and accelerate lineage sorting. Both mechanisms operate independently of ecological adaptation, challenging the long standing assumption ^39,41,42^ that neutral processes are ineffective in large bacterial populations. The simulations are necessarily simplified (e.g., 10 loci, no complex selective landscapes), but the qualitative outcome that neutral perturbations generate rapid diversification is robust across tested parameters.

### Reconciling genetic monomorphism and ancestral diversity

A paradox of the *Sulfitobacter* radiation is the extreme genetic monomorphism within each clade (within clade ANI > 99.64%) alongside the high ancestral polymorphism inferred from the large ancestral population size and extensive phylogenetic discordance. Three factors resolve this apparent contradiction. First, the recency of divergence (approximately 20,000 years) limits the time available for mutation accumulation within clades. Second, although direct measurement of within-clade recombination is precluded by the lack of polymorphic sites, any existing recombination would act as a cohesive force homogenizing genomes. Third, minimal cross-clade gene flow prevents allele introgression from other clades. This scenario contrasts with ecotype models, where periodic selection or genetic drift purges diversity ^43^. In *Sulfitobacter*, neutral processes dominate, with recombination acting as both a diversifying force (preserving ancestral polymorphisms) and a cohesive force (maintaining within-clade uniformity).

## Conclusion and future directions

This work bridges eukaryotic and prokaryotic speciation paradigms, demonstrating that ILS can drive rapid, neutral diversification in bacteria. By decoupling speciation from selection, these findings expand the framework for understanding microbial diversity, emphasizing the role of demographic history alongside classical adaptive models. As the first documented case of prokaryotic ILS, *Sulfitobacter* opens the question of how widespread this phenomenon is among bacteria. Comparative studies in taxa with high recombination rates, such as *Vibrio splendidu*s, *Streptococcus mutans*, *Pelagibacter ubique*, and *Helicobacter pylori* ^44–46^, could reveal whether ILS is rare or common. More broadly, these results suggest that neutral processes should be considered alongside selection in models of bacterial speciation, refining our understanding of microbial species boundaries and the evolutionary forces shaping global biodiversity.

## Methods

### Sample collection and bacterial isolation

We analyzed 74 *Sulfitobacter* strains collected from 16 oceanic regions and 15 ecosystem types (Fig. 1, Table S1), including 70 newly isolated genomes from diverse marine environments and four public genomes (Table S3). The 70 new isolates were obtained by multiple research teams as described below.

#### Marine isolates from the Marine Culture Collection of China (contributed by Z.Z. Shao)

The 36 strains provided by Z.Z. Shao are ecologically and geographically diverse, encompassing a wide range of marine environments and hosts, including deep-sea sediments (n=20), deep seawater (n=9), photic zone seawater (n=2), hydrothermal plume (n=1), coral sand (n=1), coral hosts (n = 3). Sampling sites spanned the Pacific, Atlantic, Indian Oceans, and the South China Sea. For environmental samples, isolation procedures were conducted aboard research vessels immediately after sampling. Samples were serially diluted in sterile seawater and plated on two different media: marine agar 2216 (MA; Difco) or Reasoner’s 2A (R2A) agar medium.

Incubation conditions varied according to sample type and cruise specifications (detailed in Table S1). Individual colonies were randomly selected and purified through successive streaking on fresh media plates. For coral-associated isolates (n=3), two strains (1A10444 and 1A10445) were obtained from *Gorgonacea* sp. collected from the South Atlantic Ocean, while strain M21595 was isolated from *Victorgorgia* sp. from the West Pacific Philippine Sea Basin. Coral tissue processing followed a standardized protocol: coral fragments (2-5 cm length) were thoroughly rinsed with sterile artificial seawater (ASW) and homogenized using a sterile mortar and pestle with 10 mL sterile ASW. The homogenization process was repeated three times to obtain a final volume of 3.0 mL homogenate. Serial dilutions of the homogenate in ASW were plated on Marine Agar and incubated at 28°C for 24 hours.

#### SpongeDassociated isolates (contributed by C.R. Voolstra)

Three specimens of *Echinochalina* sp. and three of *Stylissa carteri* were collected from Inner Fsar of the Central Red Sea (22°13’58.4” N, 39°01’45.6” E) at diving depths of 8-12 m, with specimens at least 5 m apart. Specimens were kept cool and transported to the laboratory in plastic bags with seawater. For each taxon, 1 cm^3^ pieces from each of the three specimens were pooled, homogenized with a sterile mortar in 10 mL sterile seawater. The supernatant was diluted 1:10 and 1:100, and 100 μL f of each was heated to 95°C for 10 minutes in a thermocycler. 100 μL of the heated and unheated suspensions were plated on three different agars (Difco, MI, ISP2) each containing 100 μg/mL cyclohexmide (to inhibit eukaryotic microorganisms), and half of the plates for each temperature treatment also contained 25 μg/mL nalidixic acid (to inhibit fast-growing gram-negative bacteria). Plates were incubated at 30°C for 2–8 weeks. Isolates were re-plated on the respective treatment agars and grown for one week. DNA was extracted using the DNeasy Plant mini kit (Qiagen) with the following modification: the initial cell lysis step was preceded by three cycles of liquid nitrogen freeze-thaw and tissue was lysed using a Qiagen tissuelyser with 0.5 mm glass beads (Sigma-Aldrich).

#### DiatomDassociated isolates (contributed by Y. Tomaru)

The axenic diatom strain *Chaetoceros tenuissimus* NIES-3715 was inoculated with water containing bacteria and a diatom virus CtenRNAV collected from the surface of Ariake Sound in western Japan in June 2004 ^47^. Although most diatom cells lysed due to virus infection, survivors grew ^48^. The resulting culture, in which diatom, virus, and bacteria coexisted, was defined as virus-resistant *C. tenuissimus* culture. The bacteria likely mediated the resistance of diatoms to viral infection ^48^. The filtrate of the culture (passed through a 0.8 µm filter) was cultured on ST agar (0.5% [wt/vol] tryptone, 0.05% [wt/vol] yeast extracts and 1% [wt/vol] agar in 0.2 µm-filtered seawater collected at Hiroshima Bay) or on Marine Agar (Difco™ Marine Agar 2216). After incubation for one week at 15 °C, four independent colonies from each plate were transferred to a new plate and used as isolates. These isolates were inoculated into 2mL of a culture medium (Marine Broth, 3.74 % w/v) (Difco™ Marine Broth 2216) and incubated at 25°C for 24h. Cells were collected by centrifugation at 17,400 × g for 5 min at 4°C. Genomic DNA was extracted using the DNeasy Blood and Tissue Kit (Qiagen) according to the manufacturer’s instructions and stored at −80°C.

#### Sporadic isolates from other research groups

Two strains contributed by D.C. Zhang were isolated from seawater of the South China Sea by serial dilution in sterile seawater and plating on marine agar 2216 (MA; Difco), followed by incubation at 20 °C for 10 days. One strain contributed by X.-H. Zhang was isolated from seawater of the East Pacific Ocean by serial dilution in sterile seawater and plating on 2216E agar, incubated at 22°C for two days. One strain contributed by C.M. Sun was isolated from cold seep sediment of the South China Sea by serial dilution in sterile seawater and plating on 2216E agar, incubated at 28 °C for 2-3 days. One strain contributed by I. Obernosterer was isolated from a microcosm experiment containing *Aurelia aurita* dissolved organic matter, as described previously ^49^.

All purified isolates were preserved in 20% (v/v) glycerol at -80°C for long-term storage. Taxonomic identification was performed by 16S rRNA gene sequencing using universal primers 27F and 1492R. The obtained sequences were analyzed using BLAST against reference *Sulfitobacter* genomes to confirm genus assignment.

### Genome sequencing, assembly, annotation, and average nucleotide identity calculation

The 29 genomes of the sponge- and diatom-associated isolates (contributed by C.R. Voolstra and Y. Tomaru, respectively) were sequenced at the Hubbard Center for Genome Studies, University of New Hampshire using Illumina HiSeq 2500 to generate 250 bp paired-end reads. The remaining 41 genomes were sequenced at Beijing Genomics Institute (BGI, China) using MGISEQ-2000 150 bp paired-end sequencing platform. Prior to assembly, adaptors were removed from the raw reads, and low-quality ends were trimmed using Trimmomatic-0.33 ^ref50^. Contigs were assembled from high-quality paired-end reads using SPAdes v3.9 ^ref51^ with default parameters; contigs longer than 1,000 bp with k-mer coverage >5 were retained. Assembly quality was assessed with CheckM-0.9.7 ^ref52^ (Table S1). Gene prediction and open reading frame annotation were performed with Prokka-1.14 ^ref53^. Pairwise 16S rRNA gene identity was calculated using BLAST 2.6.0+ ^ref54^, and whole-genome average nucleotide identity (ANI) between pairwise strains was calculated with fastANI ^55^.

### Phylogenomic reconstruction and ortholog identification

To place the 70 new genomes in a broader phylogenetic context, we downloaded 73 published *Sulfitobacter* genomes from the NCBI genome database (22 January 2022; Table S3) and used three *Tateyamaria* genomes as outgroups ^56^. Single-copy orthologous gene families shared by all *Sulfitobacter* strains and the outgroups were identified with OrthoFinder 2.2.1 ^57^. Protein sequences of each orthologous family were aligned using MAFFT v7.480 ^ref58^ with default parameters, and unreliable alignment sites were trimmed with TrimAl v1.4 ^ref59^ in automated mode. A concatenated alignment of 201 single-copy orthologous families was used to construct a maximum-likelihood phylogeny with IQ-TREE 1.6.2 ^ref60^ under the LG+C20+G profile mixture model and 1,000 ultrafast bootstrap replicates. The tree was visualized with iTOL ^61^. The resulting phylogeny showed that the four public *Sulfitobacter* strains (R18-1, R18-2, 15WGC, and HI0054) clustered within the clades formed by the 70 new strains (Fig. S1). Pairwise ANI among all 74 strains was approximately 95% (Fig. 1), corresponding to the operational boundary for prokaryotic species ^55^. For each clade, if multiple members were sampled from identical ecosystems, geographic locations, and research cruises (Fig. 1), only one strain was retained for downstream analyses to minimize biases from clonal expansion during sample handling and cultivation. This yielded 45 non-duplicate strains (Fig. 1, Table S1).

For these 45 strains, we re ran OrthoFinder to obtain a refined set of single copy orthologous gene families. We retained 2,117 families present in all 45 strains. For each family, nucleotide alignments were generated using MAFFT v7.480 ^ref58^ guided by the corresponding amino acid alignment, followed by TrimAl trimming. Twenty eight families contained fewer than three non redundant sequences and were excluded from gene tree construction, leaving 2,089 families. For each of these, a maximum likelihood gene tree was inferred using IQ TREE with the best fit substitution model (ModelFinder ^62^) and 100 non parametric bootstrap replicates.

### Quartet analyses for phylogenetic discordance and recombination

#### Discordance among four distinct clades

All possible quartets consisting of four strains from four different clades were examined, yielding 38,568 quartets. These were grouped into 1,001 “QUARTETs” by clade composition (e.g., QUARTET_C1 C2 C9 C12; Fig. 2A). For each QUARTET, QuartetScores ^63^ counted, across the 2,089 unrooted gene trees, the number of trees supporting each of the three possible unrooted topologies. The proportion of trees supporting each topology was calculated for every QUARTET (Fig. 2B).

#### Detection of recombination between clades

To quantify recent homologous recombination, we constructed quartets comprising strains from either two or three clades (Fig. 4A). A total of 105,512 quartets were grouped into 400 QUARTETs by clade composition. A gene tree in which two strains from the same clade are separated into different branches (Fig. 4B) indicates recent recombination between clades. For each QUARTET, we calculated the proportion of gene trees showing such mixed clade membership (Fig. 4C).

### Phylo SNP analysis

For the four focal clades (C1, C2, C9, C12), we defined a phylogenetically informative SNP (phylo SNP) as a nucleotide site that is monomorphic within each clade but polymorphic between clades, with exactly two alleles across the 45 non duplicate strains. A biallelic phylo SNP can support one of seven possible grouping patterns (Fig. 3A): four patterns in which one clade carries the derived allele and the other three carry the ancestral allele (e.g., C1 specific), and three patterns in which the four clades are split into two pairs (C1+C2 vs. C9+C12; C1+C9 vs. C2+C12; C1+C12 vs. C2+C9). We scanned the alignments of all 2,117 core genes to identify all phylo SNPs and assigned each to one of the seven types.

For each orthologous gene family, we counted the number of distinct phylo SNP types present (0–7; Fig. 3B). For each pair of phylo SNPs within the same gene family that belonged to different types (i.e., supporting conflicting topologies), we calculated the distance (bp) along the alignment between the two SNP positions (including only intervals with no intervening variable sites). The distribution of these distances was summarized across all gene families (Fig. 3C).

### Demographic inference and estimating ancestral recombination rate

#### Demographic history

To quantify the demographic history of the clades C1, C2, C9, and C12, we analyzed unfolded four-dimensional site frequency spectrum (SFS) generated from the core-gene alignments using dadi.CUDA ^31^, which is extended to fit four-population demographic models. Compared to other approaches for demographic inference, including Approximate Bayesian Computation (ABC), dadi.CUDA provides a full likelihood calculation by using the complete information of SNPs, whereas ABC typically bypasses the exact likelihood calculation by relying on summary statistics that may not capture the full genetic information. Thus, dadi.CUDA ^31^ is more suitable for this study. We fitted a simple model comprising three ancestral populations and four descendant populations (Fig. 5B), and estimated the effective population sizes of all seven populations independently and split times under the assumption of constant population size. Though incorporating population size changes can avoid biases in divergence time estimation and model choice ^64^, assuming the constant population size is robust here in that our major goal is to investigate recombination and lineage sorting in the ancestral population. We ran the program multiple times with different starting values to ensure convergence. We assessed the model’s goodness-of-fit by maximizing the model likelihood and by visual inspection of the residuals between the SFS generated by the inferred model and the real data. Assuming a mutation rate of 10^-10^ substitutions per site per generation, consistent with *Sulfitobacter pontiacus* EE 36 ^32^, we obtained an ancestral population size (*N_0_*) of 2.08 × 10^8^. To estimate the absolute timescale of the radiation, we assumed a generation time of one day, as previously determined for pelagic Roseobacter ^33,34^.

#### Ancestral recombination rate

We quantified the ancestral recombination rate (*G_0_*) of the four focal clades (C1, C2, C9, C12) by performing simulations under varying *G_0_* to fit the observed decay of linkage disequilibrium (LD) with distance. LD was measured using the squared correlation coefficient (*r^2^*) between pairs of biallelic phylo-SNPs. For a pair of biallelic phylo-SNPs giving rise to four possible genotypes (00, 01, 10, and 11) with frequencies *f*_00_, *f*_01_, *f*_10_, and *f*_11_, *r^2^* is calculated as 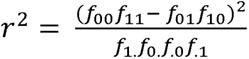, where variables with dots denote marginal probabilities, *f*_1._ = *f*_00_ + *f*_01_, *f*_.0_ = *f*_00_ + *f*_10_, *f*_1._ = *f*_10_ + *f*_11_, and *f*_.1_ = *f*_01_ + *f*_11_.

We identified 113,206 biallelic phylo-SNPs from the concatenated alignment of the four focal clades and calculated *r^2^* for all pairs of biallelic phylo-SNPs within 1 kb distance. To examine how LD decays with distance (bp), we binned the data with a non-overlapping window of 50 bp and plotted the average *r^2^* against the midpoint of each distance bin (red dots in Fig. 5A). The population mutation rate (θ*_0_*) was assumed to be consistent with the observed data (θ*_0_*=2*N*_0_μ=0.041), and *G_0_* was allowed to vary.

In prokaryotes, recombination is modelled as a gene conversion-like event with an initiation rate *g* per site per generation and the average tract length *L* ^65,66^. Because *L* is unknown but likely on the order of 1,000 bp, we first assumed *L*= 1,000 bp. The population recombination rate is *G*_0_=2*N*_0_*g*. We varied *G*_0_ from 0.02 to 0.3. For each candidate *G*_0_, we generated 10,000 random SNP patterns using msPro ^66^ under the demographic model inferred by dadi (Fig. 5B) and computed the expected LD decay (dashed lines in Fig. 5A). The best fitting *G*_0_ was selected by minimizing the sum of squared differences between the observed and simulated mean *r^2^* values across distance bins. To assess robustness, we repeated the fitting procedure with different assumed tract lengths *L* (e.g., 500 bp and 2,000 bp; Fig. S5). The recombination to mutation rate ratio was then calculated as *G*_0_/θ*_0_*.

### Neutral simulations of rapid diversification

We performed individual based simulations to test whether neutral processes alone could generate rapid diversification in the absence of adaptive selection. The simulation model was haploid with non overlapping generations. A large ancestral population was modelled as *K* subpopulations using a stepping-stone model, with migration allowed only between neighboring subpopulations (Fig. S6), thereby approximating a single panmictic population when migration is frequent. Each subpopulation comprised *N* haploid individuals. Each individual carried a haplotype composed of *L* incompatibility loci, with ancestral and derived states denoted as 0 and 1, respectively. Recurrent mutations between the two allelic states occurred at a rate μ per locus per generation. Homologous recombination replaced a contiguous block of loci from a donor genome into a recipient genome, converting the recipient into a recombinant. In each subpopulation, recombinants arose with a probability *p* per generation, and the number of loci transferred in a single recombination event was drawn from a Poisson distribution with a mean value of *r*. Genetic incompatibility was quantified as the immediate survival rate of a recombinant (ω), which decreases as the number of incompatible loci (*d*) between the recipient and the donor increases. We modeled this with a linear function, ω = 1 − *ds*, where *s* is the strength of purifying selection acting to eliminate an incompatible locus.

The simulation integrated recombination, mutation, and migration, updating the population generation by generation as follows. For each subpopulation, each individual in the offspring generation was assigned as a recombinant with probability *p* or as a non-recombinant with probability 1-*p*. Non-recombinant offspring were generated by randomly sampling and copying a single parental individual. Recombinant offspring were generated by randomly selecting two parents, transferring a continuous block of loci from the donor to the recipient, and accepting the resulting recombinant with probability ω (the survival rate). This process was repeated until *N* successful individuals had been produced to constitute the offspring generation. Mutations were introduced at rate μ per generation per locus. Finally, migration was restricted to neighboring subpopulations (e.g., subpopulations 1 and 3 are neighbors of subpopulation 2) and occurred at a rate *m* per generation.

We simulated three evolutionary scenarios under this neutral model (i.e., without adaptive selection). The first scenario was the null model, which used the following baseline parameters: the number of subpopulations *K* = 10; haploid individuals per subpopulation *N* = 1,000; number of loci *L* = 10; recombination probability *p* = 0.05; mean number of loci transferred per recombination event *r* = 1; mutation rate μ = 10^-6^; strength of purifying selection *s* = 0.2; migration rate *m* = 10^-5^ per generation. The simulation duration was 2 × 10^7^ generations. The second scenario was a mutation surge: all parameters were the same as the null model, except that μ was increased 100 fold (to 10^-4^) during a 5 × 10^6^ generation perturbation window in the middle of the simulation (from *t* = 1.0 × 10^7^ to 1.5 × 10^7^ generations). The third involves a population bottleneck: all parameters were the same as the null model, except that *N* was reduced to one tenth of its ancestral value (to 100 individuals) during the same 5 × 10^6^ generation perturbation window. After the perturbation windows, the simulation continued under the baseline parameters for the remainder of the total 2 × 10^7^ generations. For each scenario, we recorded the haplotype composition of each subpopulation after every generation. We considered a new lineage (or clade, shown in a new color in Fig. 6 and Fig. S6) when a novel haplotype spread and reached a frequency >90% in a subpopulation.

### Identification of potential genetic barriers to gene flow

We searched for the CRISPR-Cas systems and the Restriction-Modification (R-M) systems, both of which have been proposed to influence the recombination frequency between bacteria ^67–71^. CRISPR-Cas systems were identified using CRISPRCasFinder ^72^ with default parameters; only CRISPR arrays with an evidence level greater than one were retained. R-M systems were identified by searching the predicted protein sequences of all 45 genomes against the REBASE database ^73^ using DIAMOND ^74^; multiple hits with an E-value < 1e-5 for each query were manually inspected according to R-M system classification and genomic positions. For type II and type III R–M systems, genes were assigned to the same system when they matched the corresponding restriction and methylation subunits of a given R–M system ^75^, and were separated by fewer than five genes. Type I R–M systems were identified using the same criterion, except that an additional “specificity” subunit was required ^75^. Type IIG is a subtype of type II R-M system, in which the restriction and methylation subunits are encoded by a single gene; thus, only one gene was identified as type IIG R-M system if it matched such a type. Type IV R-M systems, which comprise one or two restriction subunits only ^75^, were assigned when the best hit corresponded to a one-subunit type IV system. This procedure was performed for all members of all 14 clades.

## Data availability

The raw reads and assemblies for the 70 new strains used in this project have been uploaded to the NCBI under accession number PRJNA628848.

## Code availability

All scripts used for the demographic inference are available at https://github.com/keigouematsu?tab=repositories; all scripts for gene tree construction, quartet analyses, R-M systems identification, and simulation of the neutral speciation are available at https://github.com/luolab-cuhk/Sulfito-BSC.

## Acknowledgments

HL is supported by the Hong Kong Research Grants Council General Research Fund (14107625) and the Shenzhen Science and Technology Committee (JCYJ20180508161811899). XW is supported by the National Natural Science Foundation of China (42406091).

## Author Contributions

X.W. and H.L. conceived the study; X.W., H.I. and H.L. designed research; X.W. performed the bioinformatics, K.U. performed the LD analysis, K.U. and T.A. performed the demographic inference, and H.N. performed the simulation of speciation burst under neutral scenarios. Z.S. contributed 36 isolates collected from global ocean sites; C.R.V. contributed the sponge-associated isolates from the Red Sea; K.K. and Y.T. contributed the diatom-associated isolates from the Japanese Sea; X.Z., D.Z, C.S., and I.O. contributed 2, 1, 1, and 1 isolates from unique ecosystems and diverse regions, respectively; G.L. and X.L. handled the isolates in the expanded datasets and performed the whole genome sequencing; X.W., K.U., Y.T., C.R.V., H.I. and H.L. wrote the paper. C.R.V. provided comments at various stages of the work.

## Figure legends

**Fig. S1. Expanded phylogenomic tree of *Sulfitobacter* including public genomes.** The IQ-TREE maximum-likelihood phylogeny based on 201 single-copy orthologous gene families shared by 70 *Sulfitobacter* genomes contributed by the present study and 73 publicly available *Sulfitobacter* genomes downloaded in January 2022. Solid circles at the nodes indicate the frequency of the group defined by that node is at least 90% in the 1,000 ultrafast bootstrapped replicates. The clades C1, C2, C9, and C12 together comprise 63 genomes, which are highlighted in different colors.

**Fig. S2. Sliding**D**window nucleotide identity between clades along the core genome.** The distribution of between-clade nucleotide identity along the core genome of the four focal clades (C1, C2, C9, and C12). The concatenated core genome alignment was used to calculate the mean nucleotide identity within 1,000 bp sliding windows at a step of 500 bp. The red horizontal dashed line sets the 99% nucleotide identity.

**Fig. S3. Distribution of restriction**D**modification (R-M) systems across clades across the 45 strains and absence of CRISPR**D**Cas systems.** (Left) The rooted maximum-likelihood phylogenomic tree where clades (C1, C2, C9, and C12) with at least three members are colored. (Right) The copy number of R-M systems in each genome. The type I to IV R-M systems are marked in yellow, purple, green, and orange, respectively. No CRISPR Cas system was detected in any sequenced genome.

**Fig. S4. Between-clade nucleotide sequence identity of clade-subset gene families and background single-copy orthologs (SCOs) in *Sulfitobacter*.** Violin plots show pairwise between-clade nucleotide identity for clade-subset genes (red) and background SCOs (blue) across six clade pairs. For genes shared by two clades, identity was calculated for that clade pair; for genes exclusive to one clade, identity was calculated among the other three clades. *P* values were calculated using one-sided Mann-Whitney U tests (subset > background). Panel titles indicate the subset categories contributing to each clade pair.

**Fig. S5. Linkage disequilibrium (LD) decay with varying recombination tract lengths.** The decay of LD of the four focal clades (C1, C2, C9, and C12) with varying tract lengths. The horizontal and vertical axes represent the distance (bp) between pairwise biallelic phylo-SNPs and the *r^2^* measure, respectively. The horizontal dashed line marks the expected LD under the inferred demography when the distance of pairwise biallelic phylo-SNPs is ∞. Distances are binned with a window size of 50 bp, and the red dots denote the observations from the *Sulfitobacter* population under study. The dashed curves represent the expected decay of LD under different tract lengths and a fixed population recombination rate (*G* = 0.1), assuming the demographic model shown in Fig. 5B.

**Fig. S6. Null simulation with no perturbation.** Representative result from simulation under the null model, where *K* = 10, *N* = 1000, *L* = 10, *p* = 0.05, *r* = 1, μ = 10^-6^, *s* = 0.2, and *m* = 10^-5^ are assumed. In the initial state, all individuals were assumed to have the same haplotype with the ancestral allele 0 at all loci (shown in magenta). New haplotypes are shown in different colors. The proportion of the haplotypes is monitored and plotted for each subpopulation.

