## Supplementary figures for "Rapid Speciation Characterized by Incomplete Lineage Sorting in the Globally Distributed Bacterium *Sulfitobacter*"

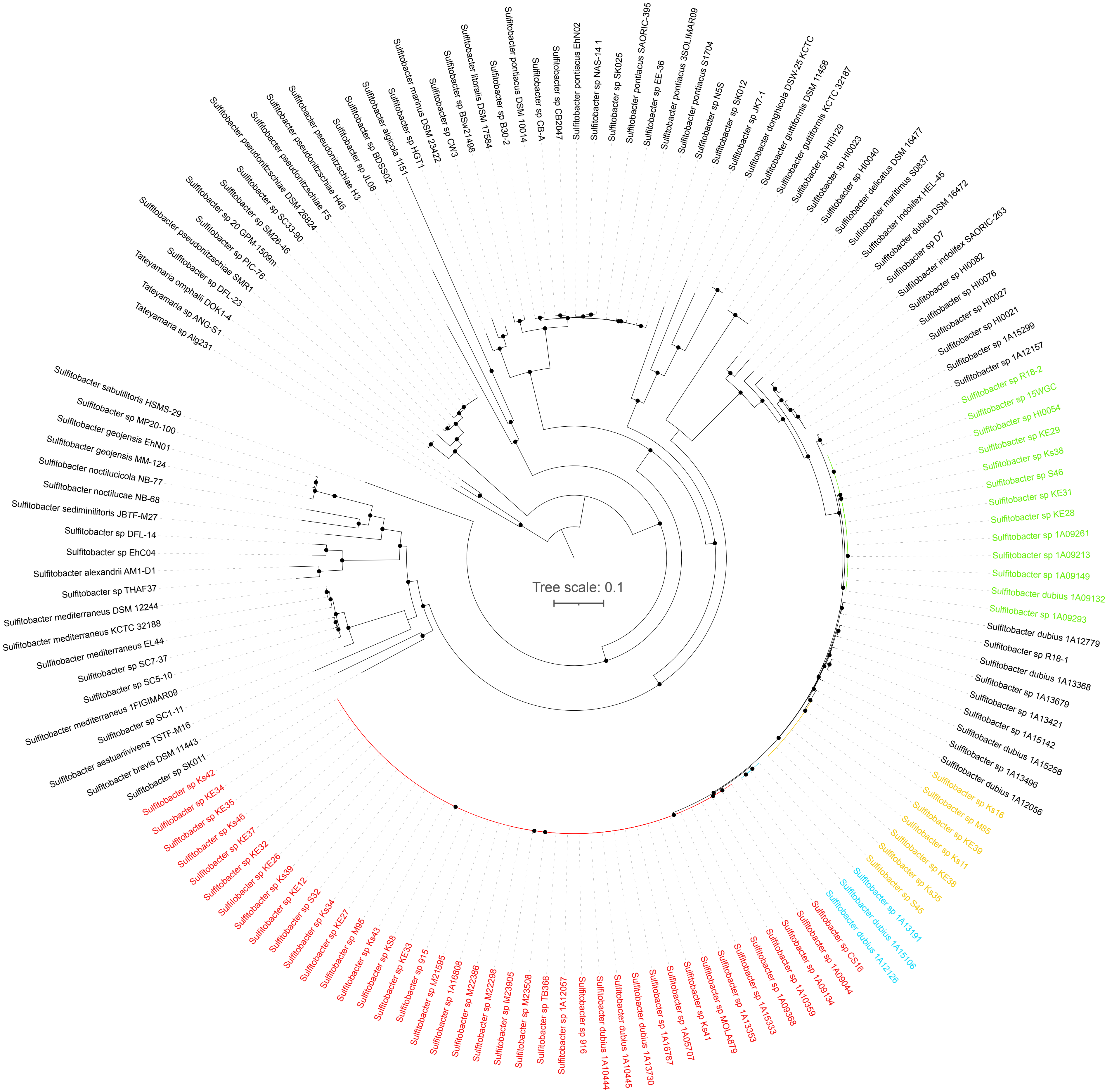


**Fig. S1. Expanded phylogenomic tree of *Sulfitobacter* including public genomes.** The IQ-TREE maximum-likelihood phylogeny based on 201 single-copy orthologous gene families shared by 70 *Sulfitobacter* genomes contributed by the present study and 73 publicly available *Sulfitobacter* genomes downloaded in January 2022. Solid circles at the nodes indicate the frequency of the group defined by that node is at least 90% in the 1,000 ultrafast bootstrapped replicates. The clades C1, C2, C9, and C12 together comprise 63 genomes, which are highlighted in different colors.

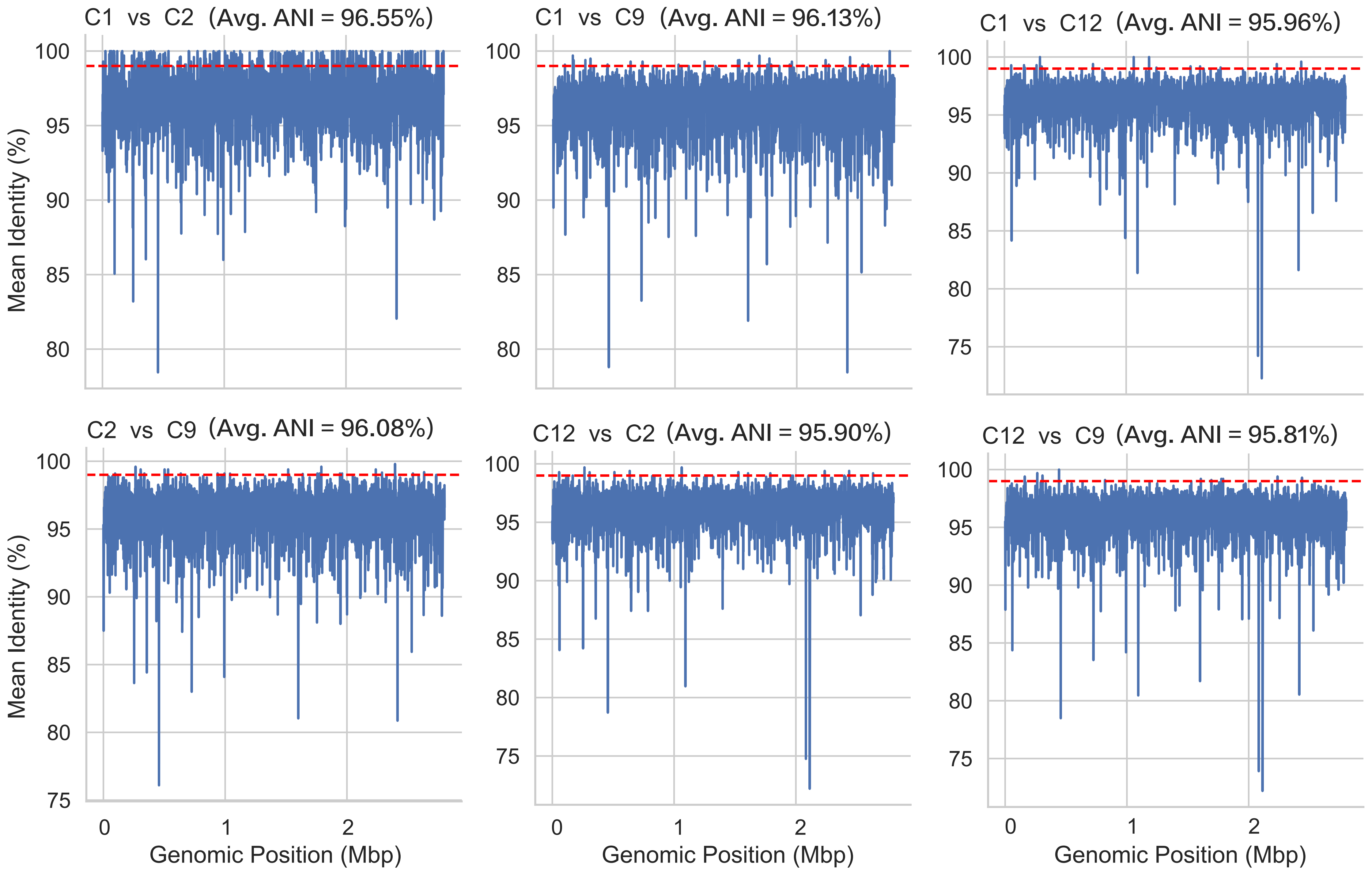


**Fig. S2. Sliding-window nucleotide identity between clades along the core genome.** The distribution of between-clade nucleotide identity along the core genome of the four focal clades (C1, C2, C9, and C12). The concatenated core genome alignment was used to calculate the mean nucleotide identity within 1,000 bp sliding windows at a step of 500 bp. The red horizontal dashed line sets the 99% nucleotide identity.

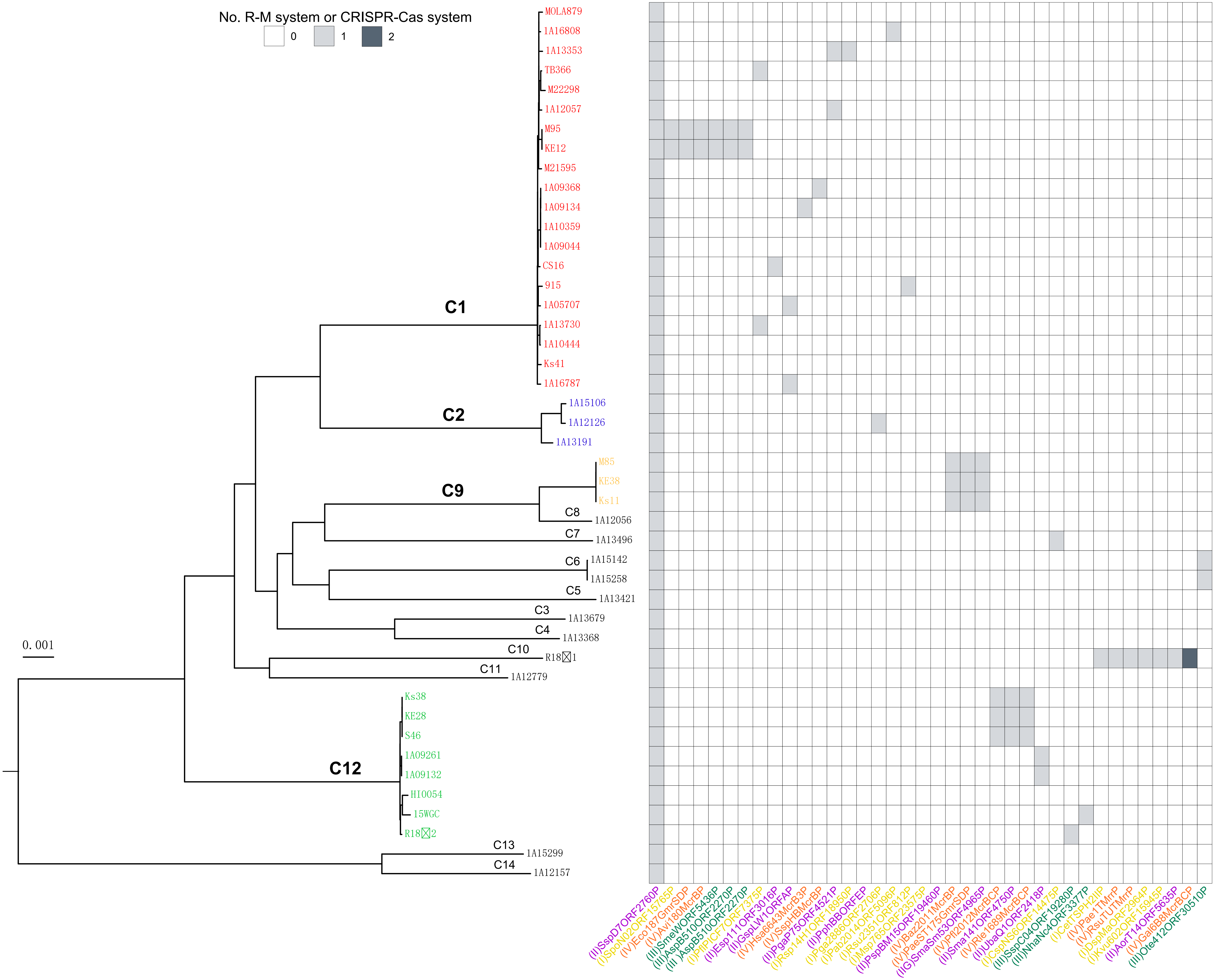


**Fig. S3. Distribution of restriction-modification (R-M) systems across clades across the 45 strains and absence of CRISPR-Cas systems.** (Left) The rooted maximum-likelihood phylogenomic tree where clades (C1, C2, C9, and C12) with at least three members are colored. (Right) The copy number of R-M systems in each genome. The type I to IV R-M systems are mark**ed** in yellow, purple, green, and orange, respectively. No CRISPR-Cas system was detected in any sequenced genome.

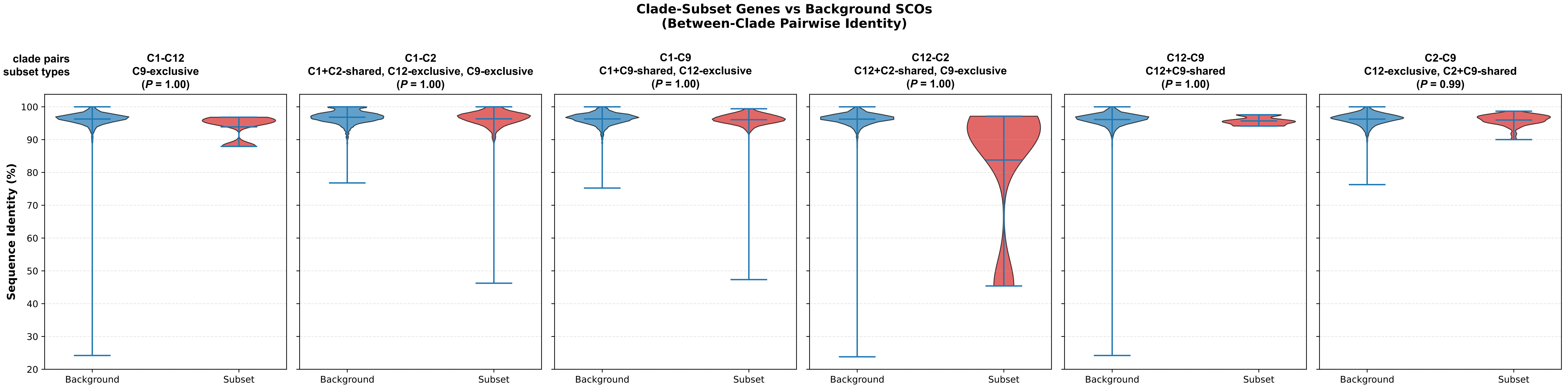


**Fig. S4. Between-clade nucleotide sequence identity of clade-subset gene families and background single-copy orthologs (SCOs) in *Sulfitobacter*.** Violin plots show pairwise between-clade nucleotide identity for clade-subset genes (red) and background SCOs (blue) across six clade pairs. For genes shared by two clades, identity was calculated for that clade pair; for genes exclusive to one clade, identity was calculated among the other three clades. *P* values were calculated using one-sided Mann-Whitney U tests (subset > background). Panel titles indicate the subset categories contributing to each clade pair.

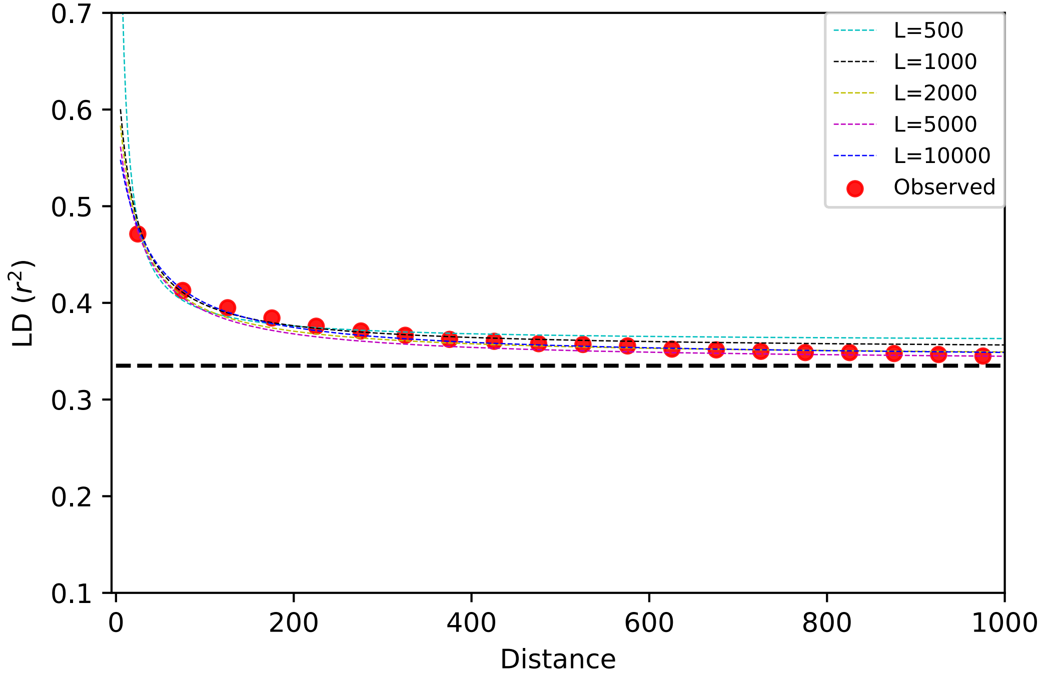


**Fig. S5. Linkage disequilibrium (LD) decay with varying recombination tract lengths.** The decay of LD of the four focal clades (C1, C2, C9, and C12) with varying tract lengths. The horizontal and vertical axes represent the distance (bp) between pairwise BiPs and the $r^2$ measure, respectively. The horizontal dashed line marks the expected LD under the inferred demography when the distance of pairwise BiPs is ∞. Distances are binned with a window size of 50 bp, and the red dots denote the observations from the *Sulfitobacter* population under study. The dashed curves represent the expected decay of LD with different tract lengths and a fixed population recombination rate ($G$ = 0.1), calculated by assuming the estimated demographic model shown in Fig. 5B.

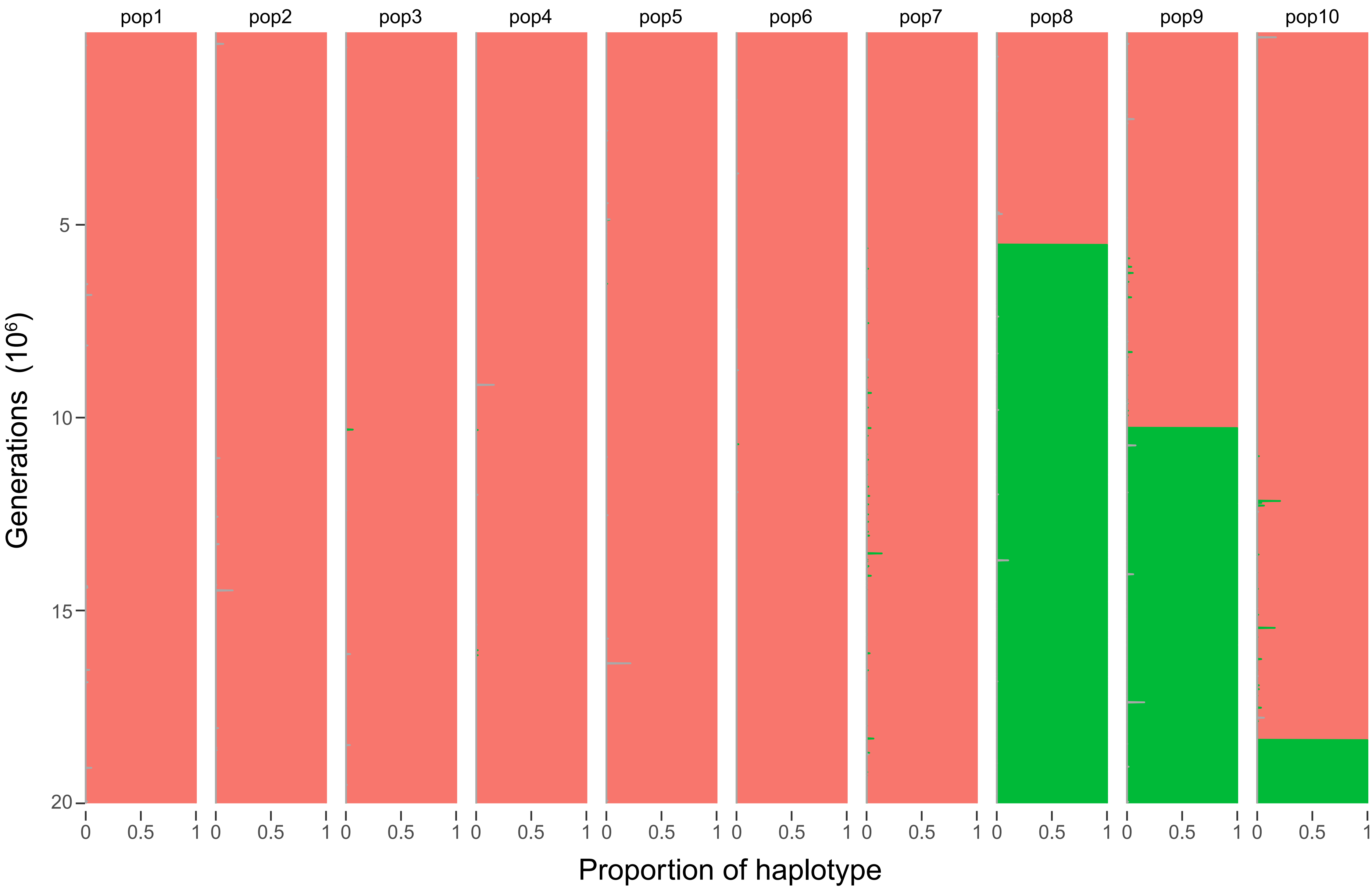


**Fig. S6. Null simulation with no perturbation.** Representative result from simulation under the null model, where $K$ = 10, $N$ = 1000, $L$ = 10, $p$ = 0.05, $r$ = 1, $\mu = 10^{-6}$, $s$ = 0.2, and $m = 10^{-5}$ are assumed. In the initial state, all individuals were assumed to have the same haplotype with the ancestral allele 0 at all loci (shown in magenta). New haplotypes are shown in different colors. The proportion of the haplotypes is monitored and plotted for each subpopulation.
