## Supplementary information for "Rapid Speciation Characterized by Incomplete Lineage Sorting in the Globally Distributed Bacterium *Sulfitobacter*"

**This file includes the following items:**

Supplementary Results

References

Supplementary Results

1.1 Exploring potential recombination barriers of the Sulfitobacter clades

Given the very high recombination rate in the ancestral population, effective recombination barriers must have been established to initiate speciation. Such barriers may arise through genetic changes, ecological adaptation, and physical or geographic separation ^1–4^. Geographic separation, however, is unlikely to be a speciation driver, because the 74 *Sulfitobacter* members were recovered from widely distributed locations, and multi-member clades like C1, C2, C9, and C12, each comprising strains from multiple oceanic regions (Fig. 1). Additionally, host preference does not explain speciation, as members from the same clade are associated with multiple host lineages such as corals and sponges (Fig. 1).

To explore potential ecological factors other than host selection, we examined two types of gene families: i) clade-specific gene families that are universally and exclusively present in one clade but absent in the remaining clades; ii) clade-excluded gene families that are absent in a specific clade but present in the rest. These gene families may provide some clues to the functional differentiation between clades. Focusing on the four multi-member clades (C1, C2, C9, and C12), we identified 163 orthologous gene families (Table S4) potentially contributing to ecological divergence on the basis of presence-absence patterns using OrthoFinder 2.2.1 ^ref5^. Of these, 102 lacked functional annotation. The remaining 61 gene families were assigned KEGG annotations using one representative member from each orthologous gene family. Although these gene families spanned diverse functional categories, including transport and uptake systems, metabolic and enzymatic pathways, and regulatory elements, most comprised only one or a few components of otherwise incomplete pathways. As a result, these data alone provided only limited support for clear ecological niche differentiation.

Even so, several patterns suggested possible metabolic specialization among clades. The clearest signal was observed in Clade C9, which specifically possesses a set of functionally related genes involved in polyhydroxyalkanoate (PHA) synthesis, including *phaC*, *croR*, and components of the ETF electron transfer system. The co-occurrence of multiple genes within a single carbon storage pathway suggested that C9 may be adapted to environments with fluctuating carbon availability, where PHA accumulation may confer a competitive advantage. Other clades showed more limited but potentially informative signals.Clade C1 carried a clade-specific glutamine synthetase gene family, whereas Clade C2 contained a sulfate permease gene family in Clade C2 and a *LacI*/*GntR*-type transcriptional regulator in C2. Clade C12 carried specific gene families involved in xanthine degradation (*xdhA*/*xdhC*), pentose and glucuronate interconversion (*uxaC*/*uxuB*), and organic nitrogen acquisition (spermidine/putrescine transporter, urea carboxylase). Notably, C12 also lacked orthologous gene families related to two peptide/nickel ABC transporter systems and one iron-complex uptake ABC transporter cluster that were present in other clades.

A complementary KO-based analysis, in which copy numbers of each KEGG identifier were counted directly for each genome, revealed that most of the differences described above reflect quantitative copy number variation rather than strict binary presence/absence (Table S5). The main exceptions include genes corresponding to urea carboxylase and glucuronate isomerase (*uxaC*), which were specific to C12. Notably, the gene-content differences could also reflect independent post-divergence acquisition or differential retention of accessory genes via horizontal gene transfer (HGT) following an initial ILS-driven divergence, and therefore may not represent the ecological factors that established the initial barriers to recombination. Accordingly, the observed gene-content differences are insufficient, on their own, to identify ecological selection as the primary driver of divergence.

1.2 The simulated speciation process under a neutral scenario with constant evolutionary parameters

We demonstrated that speciation can occur under this neutral model in the absence of adaptive selection, where incompatibility loci initiate speciation. For instance, Fig. S6 shows the result of a single simulation run under the null model. The simulation began with all individuals having the ancestral haplotype at all loci (haplotype=0000000000, in magenta). Over time, a new haplotype arose in subpopulation 8 (shown in green) at around *t* = 6 × 10^6^ generations and became fixed. The new haplotype then migrated to the adjacent downstream subpopulation 9, where it also became fixed, and subsequently to subpopulation 10. The new haplotype also migrated to the upstream subpopulation 7, but did not reach fixation there. Across a broad range of parameter combinations, the simulations consistently produced speciation under this neutral model, and the inferred rate of speciation was qualitatively robust to parameter variation (data not shown).
